# Coral reef fish resilience and recovery following major environmental disturbances caused by cyclones and coral bleaching: A case study at Lizard Island

**DOI:** 10.1101/2024.01.27.577527

**Authors:** Simon A. Lévy, Letizia Pessina, Redouan Bshary, Zegni Triki

**Author notes:** Correspondence to: Zegni Triki.

## Abstract

Coral reef fish communities can be affected by natural disturbances such as cyclones and coral bleaching. It is not yet understood how long it takes these communities to recover from such extreme events, particularly when they occur repeatedly. To investigate this, we conducted fish surveys repeatedly between 2011 and 2022 at Lizard Island on the Great Barrier Reef in Australia. We focused on two reef sites, Mermaid Cove and Northern Horseshoe, both of which were damaged by a large-scale coral bleaching event in 2016 and 2017, as well as two cyclones that occurred in 2014 and 2015 (the cyclones hit Mermaid Cove but not Northern Horseshoe). Between 2016 and 2017, both reef sites saw a decrease in the total fish abundance of about 68 % and across most functional groups (carnivores, corallivores, herbivores, and omnivores). Despite the two sites showing different decline and recovery patterns, they both showed an improvement in fish abundance and across the majority of functional groups at both sites by 2022. The recovery reached similar numbers as those documented in the fish census data collected before the disturbances occurred. Our findings provide a case study highlighting how fish community resilience can vary on small local scales, with potential recovery if conditions are favourable over several years.

## Introduction

Although coral reefs cover a limited surface area, they are crucial ecosystems that harbour over 25 % of all fish species. Seventy-five per cent of these fish rely on corals for food, shelter, and settlement (Coker et al., 2014; Jones et al., 2004; Tarbuk et al., 1992; Spalding & Grenfell, 1997). According to estimates, the extinction of all coral reefs could lead to the loss of around 50 % of fish species due to their association with the reefs (Strona et al., 2021b). Extreme climatic events are threatening coral reefs (Hughes et al., 2017), and as a consequence, they also threaten coral reef fish communities in terms of density, diversity, and functionality (Jones et al., 2004; Pratchett et al., 2011; Richardson et al., 2018; Triki & Bshary, 2019; Wilson et al., 2006). One of the major threats to coral reefs is the prolonged El-Niño, a natural climatic event that brings warm water towards the Indo-Pacific Ocean. This increase in sea temperature can over-stress the corals and disrupt their symbiotic relationship with the photosynthetic Zooxanthella, which provides the corals with their pigmentation. When the Zooxanthella are expelled, they cause the corals to become bleached (Cai et al., 2014; Hoegh-Guldberg & Ridgway, 2016). If corals are not recolonised by Zooxanthella, they are predicted to die within approximately six months (McCook, 2002). In addition to extreme climatic events causing bleaching, cyclones are also an imminent threat to corals and can cause significant coral cover loss in exposed areas (Cheal et al., 2017; Dixon et al., 2022).

Depending on the type, frequency, and intensity of disturbances, the extent of coral loss and resulting impacts on coral reef fish communities may vary (Jones et al., 2004; Pratchett et al., 2011; Wilson et al., 2006). In the short term, biological disturbances (e.g. coral bleaching) reduce the live coral cover while leaving the structural complexity of the affected location intact since the underlying skeletons of the corals are unaffected by this type of disturbance. In contrast, physical disturbances (e.g. cyclones) immediately reduce the live coral cover and the structural complexity of the affected location (Wilson et al., 2006; Pratchett et al., 2011). Therefore, it appears that in the short term, physical disturbances have a greater impact on fish biodiversity than biological disturbances (Pratchett et al., 2011). However, over the long term (e.g., 4-5 years), biological disturbances impact the fish similarly to physical disturbances since the structural complexity of the reef is also reduced by the erosion of the underlying coral skeletons (Pratchett et al., 2008). The intensity of the disturbances also contributes to the extent of the impact on the fish community (Pratchett et al., 2011). Large disturbances that cause extensive coral loss can significantly reduce the density and diversity of fish in the area. On the other hand, minor or moderate disturbances that result in moderate coral loss can lead to a short-term increase in the local fish diversity, potentially caused by the rise in overall heterogeneity of the habitat (Triki & Bshary, 2019; Wilson et al., 2006; Jones et al., 2004; Pratchett et al., 2011).

The health and recovery of coral reefs and their associated fish communities are influenced not only by the frequency of disturbances (Osborne et al., 2017; Ortiz et al., 2018) but also by several other factors. The complexity of coral structure and composition, geographic location, and water depth can all impact the degree of resistance of reef fish to disturbances (Graham et al., 2015; Richardson et al., 2018; Strona et al., 2021a). After environmental disturbances, the corals can take up to 15 years to recover depending on the frequency, the intensity of disturbances and the exposition of the locations (Morri et al., 2015; Ortiz et al., 2018; Osborne et al., 2017; Tebbett et al., 2022). Long-term surveys are powerful tools for identifying coral recovery stages and monitoring fish communities. For example, in Thailand, between 2013 and 2019, a study was conducted to evaluate fish recovery after a major 2010 coral bleaching event (Jaroensutasinee et al., 2020). They found that fish diversity and density increased over the years as the coral reef recovered.

A recent study at Lizard Island on the Great Barrier Reef in Australia looked into how environmental disturbances such as cyclones and coral bleaching affect fish abundance and densities of different functional groups (Triki & Bshary, 2019). Triki and Bshary looked into fish survey data collected before and after major environmental disturbances that hit Lizard Island coral reefs between 2014 and 2016. They categorised fish species into 11 functional groups based on their diet to explore density changes within each group. The numbers indicated a 68 % total decline in fish densities after the disturbance, with a notable density decrease in nine of the 11 trophic groups. The trophic group of piscivores emerged as the only group with increased density after the disturbances, likely caused by the reduction of shelters for their prey (Triki & Bshary, 2019). Here, we continued surveying the two reef locations at Lizard Island, previously studied by Triki and Bshary (2019) (Figure 1). Our aim was to monitor the resilience and recovery of coral reef fish in the years after repeated disturbances (Figure 2). In the span of three years, Lizard Island experienced three extreme environmental disruptions. In 2014 and 2015, the location was hit by cyclones Ita and Nathan, respectively (Puotinen et al., 2016; Pizarro et al., 2017). A prolonged El-Niño in February and March 2016 caused a massive coral bleaching event that also impacted the Island (Hughes et al., 2017). The 2016 coral bleaching event was devastating which resulted in 60% of the coral being bleached and a loss of 51% of coral cover (Hughes et al., 2017; Stuart-Smith et al., 2018). In 2017 and 2020, the Island experienced other coral bleaching events, but they were less severe than the 2016 bleaching and more localised (Hughes et al., 2021; Tebbett et al., 2022).

**Figure 1.**
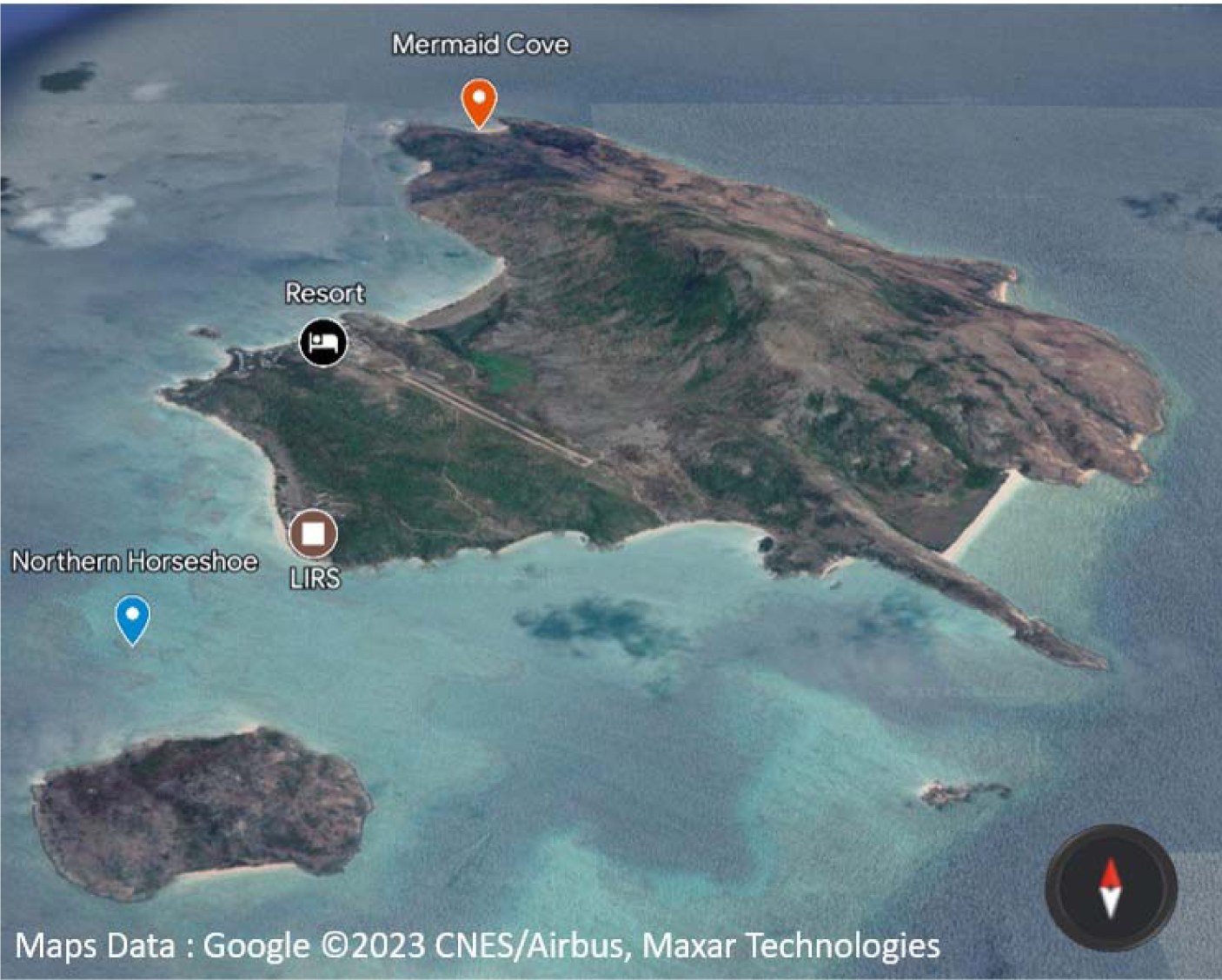
Map of Lizard Island (QLD, Australia). The map is showing the two study sites: Mermaid Cove (red) and Northern Horseshoe reef (blue). The Lizard Island research station (LIRS) and the resort are also indicated in this map.

**Figure 2.**
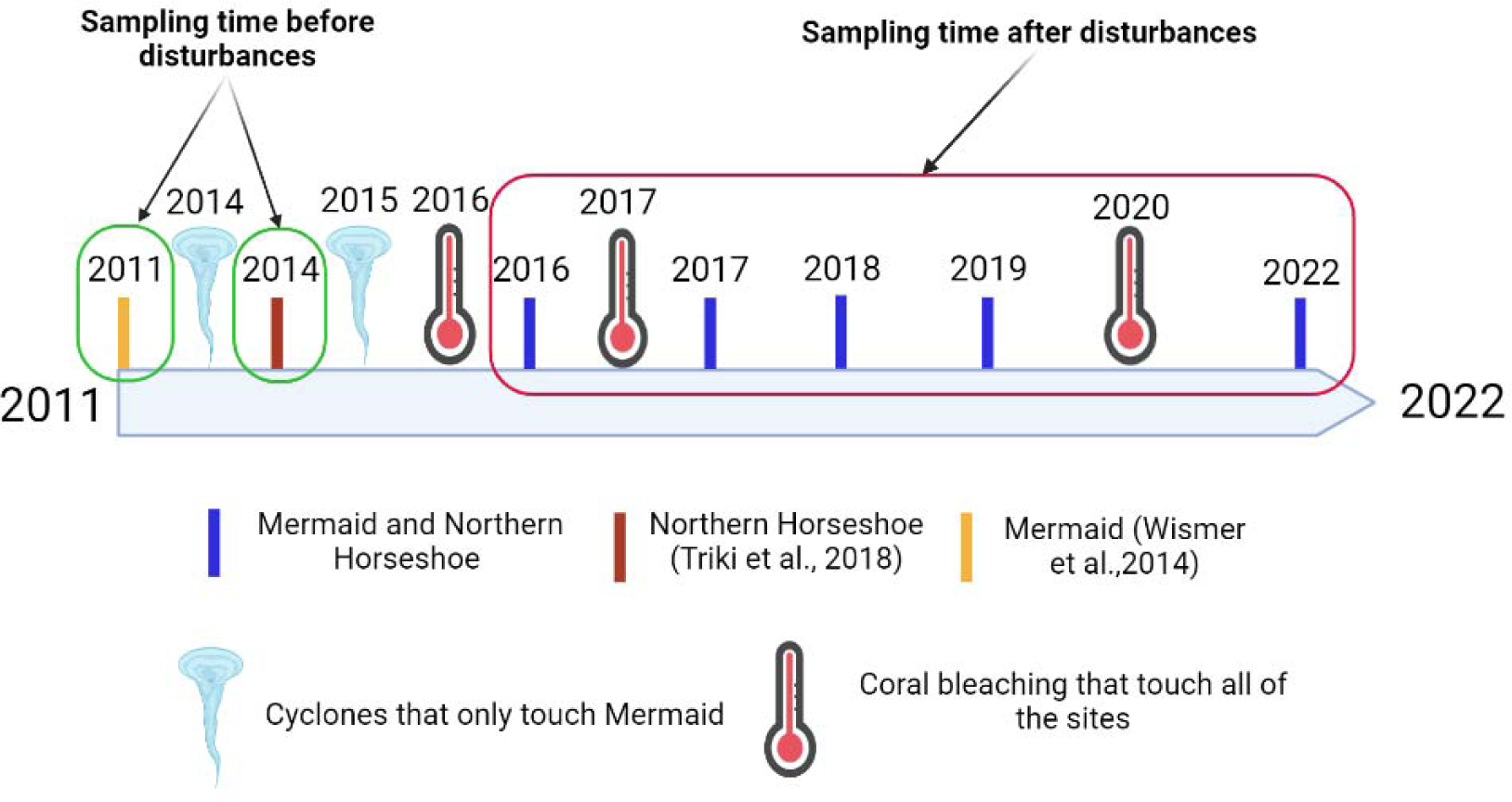
A schematic representation of the data collection timeline between the year 2011 to 2022 at the two study locations, Mermaid Cove and Northern Horseshoe.

In our study, we collected fish census data using methods similar to those used by Triki and Bshary (2019). This allowed us to track changes over time and investigate how the abundance, density, richness, and evenness of fish (that is, the abundance of each species) fluctuated before and after disturbances occurred (see timeline in Figure 2).

## Methods

### Field sites and timeline

The study was conducted on the reef around Lizard Island, Great Barrier Reef, Australia (14.6688° S, 145.4594° E). The two study locations were Mermaid Cove and Northern Horseshoe. Mermaid Cove (14.6478° S, 145.4542° E) is located in a small bay on the northern side of Lizard Island (Figure 1). It forms a continuous fringing reef of approximately 35,000m^2^ with a 1-7 meter depth. Northern Horseshoe (14.6856° S, 145.4438° E) is located on the western side of the island (Figure 1). It is a continuous reef that consists of a coral garden of approximately 17,000 m2, with a 1-4 meter depth.

We have combined our data with previously published data from Triki and Bshary (2019). The published data from the two sites, Mermaid Cove and Northern Horseshoe, were collected in 2011, 2014, 2016, and 2017. Our data collection took place between July and August in 2018 and 2019 and in December 2022 (Figure 2). Unfortunately, due to the Covid-19 pandemic lockdown, we were unable to collect data for 2020 and 2021.

### Fish census data collection

We conducted an underwater visual fish census using the same methods as Triki and Bshary (2019). In each location, the observer swam ten replicates of a 30-meter transect line on the reef. During the survey, an observer swam along each 30-meter transect line and counted the number of large visible fish, which are the species with a total body length of 10 cm or more within a 5-meter wide area. On the way back, the observer recorded the number of small visible fish, which are the species with a total body length of less than 10 cm, within a 1-meter wide area (it was done on a 2-meter wide area in 2022 unintentionally). It is important to note that only adult fish were surveyed.

The size of the species (i.e. large or small) was based on the classification by Triki et al. (2018). Each of the ten transect replicates was sampled at least 10 meters apart to minimise possible resampling of the same individuals. Fish species were identified using the Lizard Island field guide (http://lifg.australianmuseum.net.au/) and the fish guidebook by Allen et al. (2005). Overall, we identified 206 species in our surveys. All fish counts for large and small fish species were scaled per 100 m^2^.

### Fish census analyses

We estimated fish densities at each study location and period of data collection as the total number of fish per 100 m^2^. Afterwards, we categorised fish species into 11 functional groups based on their trophic level, following the methods by Triki and Bshary (2019). To simplify this step further, we opted for grouping the functional groups into four primary functional groups following the classification of Pratchett et al. (2011): (1) carnivores, which are fish that feed on other animals, this category includes invertivores (micro and macro), pisci-invertivores and piscivores, (2) corallivores, which are fish that feed on corals, (3) herbivores, which are fish that feed on plant material, this category includes browsers, grazers, detritivores, excavator and scrapers, and (4) omnivores, which are fish that feed on plant and animal material, this category includes spongivores, planktivores, and fish species with mixed diet.

Considering the various concepts and methods used to estimate diversity (Tuomisto, 2010 and 2011; Hoffmann and Hoffmann, 2008), we have focused on three aspects: (i) richness, (ii) evenness, and (iii) composition. (i) We calculated richness using the number of species per transect. (ii) We used Pielou’s evenness index to estimate the evenness of species distribution in a community. The Pielou index ranges between 0 and 1, where 1 indicates perfect evenness, and the value decreases towards 0 as the relative abundances of the species become more unevenly distributed. The Pielou index builds on the Shannon index that estimates species diversity (Pielou, 1966). We used the formula (1) to calculate first the Shannon index (H) by fitting the number of individuals for species (ni), total number of individuals (N), and number of species (S). Then, we fitted the Shannon index (H) in the formula (2) to calculate Pielou’s evenness index (E), with (S) being the number of species.

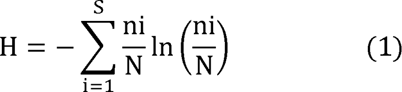

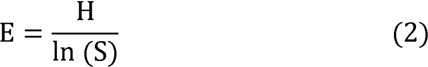

(iii) For fish composition analyses, we performed a non-metric multidimensional scaling (NMDS). The purpose of NMDS is to accurately display the position of objects (i.e., fish communities) in multidimensional space using a limited number of dimensions that can be easily visualized. This method is similar to principal component analyses, although the axes are more arbitrary. The NMDS analysis uses a strength coefficient (stress level) to estimate the goodness of fit, wherein a stress level of < 0.2 indicates certainty, < 0.1 is a good fit, and < 0.05 is an excellent fit (Yang et al. 2021).

For the diversity calculations (i.e. richness, evenness and composition), the data from 2022 were not used since the small fish number was unintentionally sampled on a 2 meter wide area instead of 1 meter.

### Statistical analyses

All data and figures were generated using the open-source software R, version 4.2.3 (R Core Team, 2022). The transect line was the statistical unit for the fish census data. All the statistical models had the year of data collection and location as predictors with a general syntax of: response.∼ year * location. We fitted General Least Square (GLS) models for fish density and Pielou’s index analyses. We transformed the data to fit normality model assumptions by using the function *boxcox()* in R language, where we applied cube root transformation on fish density data and arcsine transformation on Pielou’s index data. We used a General Linear Model (GLM) with a negative binomial distribution to analyse the richness data. For the functional group analyses, we used a set of models that were best suited to the data distribution of each functional group. That is, we fitted Linear Models (LM) for the functional groups carnivores and herbivores. Data needed log-transformation to fit the model normality assumption. For corallivores and omnivores, we fitted GLS models also with log-transformation.

For the post hoc analyses, we run *emmeans()* tests (from the package “emmeans”). Given the considerable large number of pairwise comparisons among years of data collection and locations, we opted for compact letters (a, b, c, …) display in the corresponding plots (see Figures). We provide, however, a detailed statistical table in the Supplementary Material (Table S1) with estimates, standard errors and p-values. The *emmeans()* tests take into account and correct for multiple hypothesis testing.

All fitted models met their corresponding assumptions, such as residuals’ normality and homoscedasticity. We used a set of visual plots and statistical tests, Shapiro-Wilk, Lilliefors (Kolmogorov-Smirnov) and Bartlett’s tests, to confirm that there were no violations of the assumptions.

For the composition analysis, we ran an NMDS with a Bray-Curtis dissimilarity index using the function metaMDS from the R package vegan. We selected the number of dimensions such that the stress value is equal to or lower than 0.2 (Clarke, 1993; Yan et al. 2021). For the overall comparison, we used a Permutational multivariate analysis of variance (PERMANOVA; function *adonis2()* in R language; number of permutations: 999). For the post-hoc analyses, we ran multiple pairwise comparisons with the p-value adjusted with the Holm method using the function *pairwise.adonis()* in R language. More information is available in a detailed step-by-step R code that enables the replication of the findings.

## Results

### Fish density

We found that fish density, as the number of fish per 100 m^2^, varied across the years (GLS: N = 120, X^2^ = 469.054, p < 0.001) and locations (GLS: N = 120, X^2^ = 32.724, p < 0.001). In Northern Horseshoe, the posthoc analyses revealed that fish densities significantly declined in 2017 (p < 0.001). However, in the following years, the density started returning to levels documented before the disturbances, reaching the highest values in 2022 (p < 0.001) (Figure 3A). After the disturbances, fish density in Mermaid Cove immediately declined (p < 0.001). The decline continued in 2017 (p < 0.001). Similar to Northern Horseshoe, the fish density gradually started to recover, and in 2022, it returned to the documented densities before the disturbances (Figure 3B).

**Figure 3.**
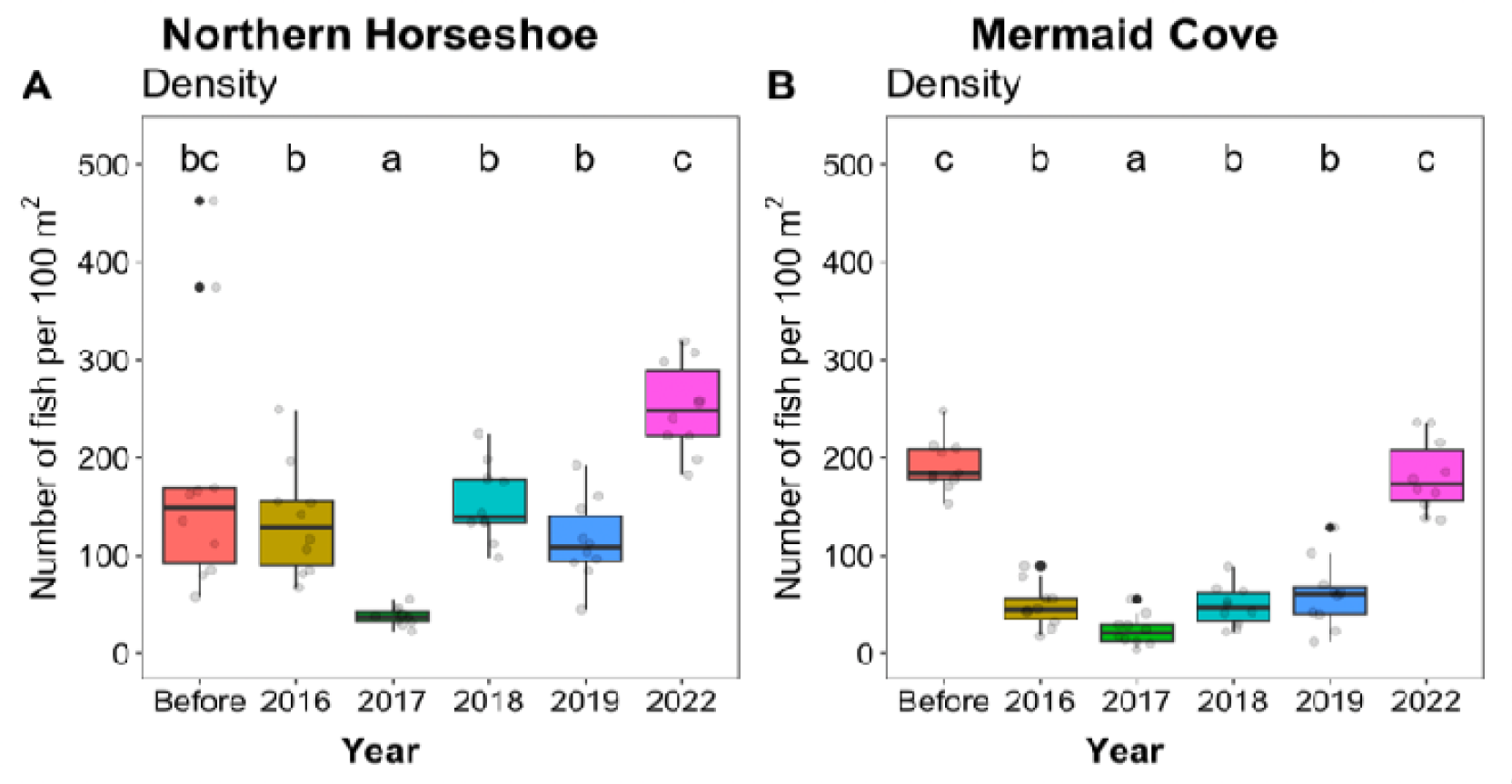
Fish density at the two study locations, Mermaid Cove and Northern Horseshoe. Boxplots of median and interquartile range of fish counts per 100 m^2^ before and in the years after the major perturbation events (see Figure 2 for timeline). Dissimilar letters above the boxplots indicate statistically significant differences (p < 0.05). The year labelled “Before” corresponds to the data from 2011 for Mermaid Cove and 2014 for Northern Horseshoe.

### Fish functional groups

All four major functional groups showed statistically significant density fluctuations across years of data collection and locations: carnivores (LM: N = 120, year, F-value = 17.821, p < 0.001; location, F-value = 4.08, p = 0.046); corallivores (GLS: N = 120, year, X^2^ = 48.101, p < 0.001; location, X^2^ = 4.174, p = 0.04); herbivores (LM: N = 120, year, F-value = 30.561, p < 0.001; location, F-value = 25.567, p < 0.001), and omnivores (GLS: N = 120, year, X^2^ = 616.96, p < 0.001; location, X^2^ = 99.029, p < 0.001). The posthoc analyses showed a sharp decline in carnivore, herbivore, and omnivore species in Northern Horseshoe in 2017, while corallivores remained unaffected (Figure 4). In the following years, especially in 2022, the analyses indicated a recovery in these functional groups (Figure 4A, E & G). After the perturbations, all four functional groups at Mermaid Cove underwent a significant decline by 2017 but eventually recovered by 2022 (Figure B, D, F & H).

**Figure 4.**
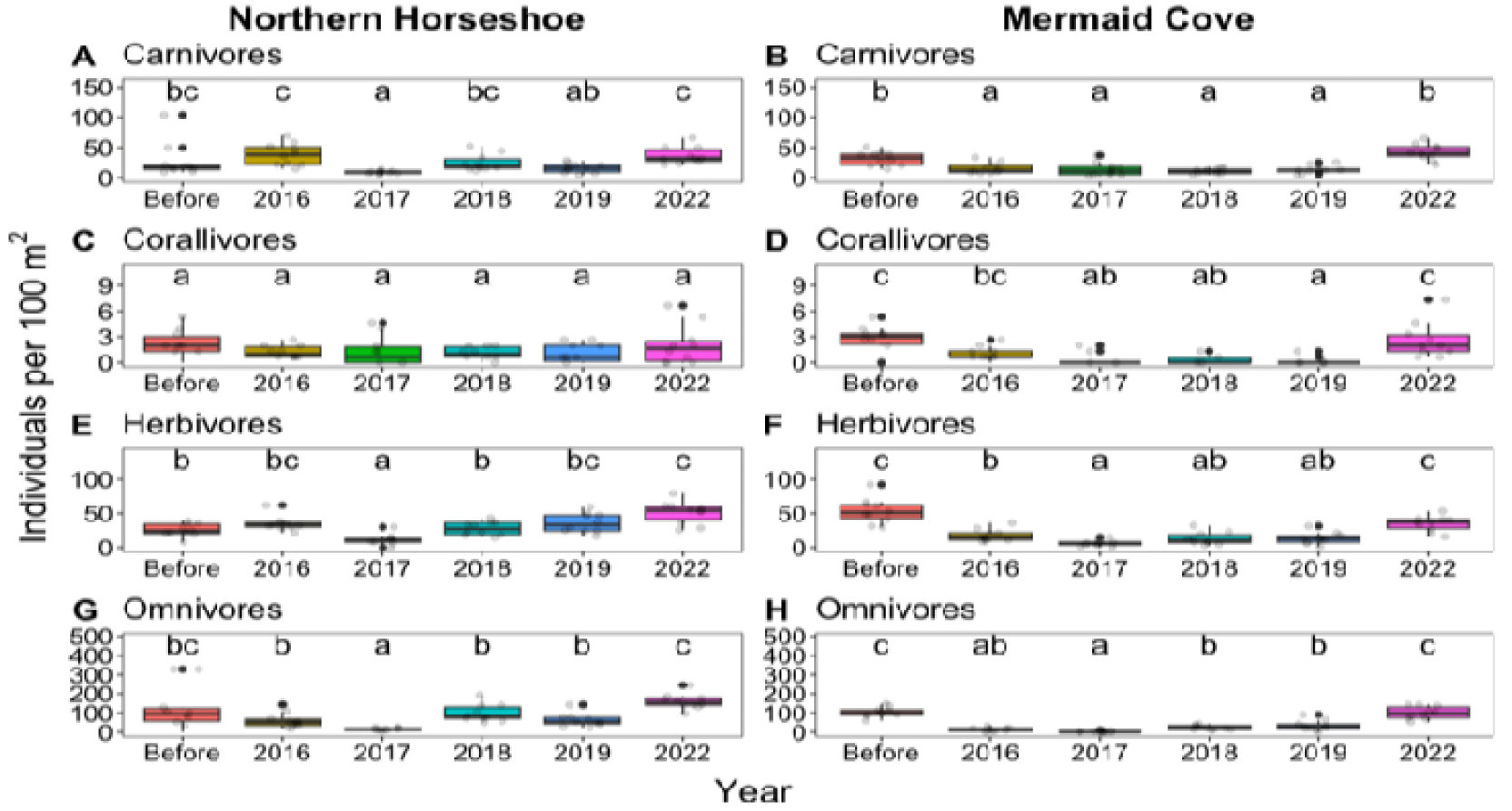
Fish density per primary functional groups at Mermaid Cove and Northern Horseshoe. Boxplots of median and interquartile range of fish densities in the functional groups (A-B) carnivores, (C-D) corallivores, (E-F) herbivores, and (G-H) omnivores. Dissimilar letters above the boxplots indicate statistically significant differences (p < 0.05). The year labelled “Before” corresponds to the data from 2011 for Mermaid Cove and 2014 for Northern Horseshoe.

### Fish Diversity

#### a. Richness

Analyses on the number of species indicated a significant variation across years of data collection (negative binomial GLM: N = 100, X^2^ = 28.248, p < 0.001) and locations (negative binomial GLM: N = 100, X^2^ = 33.830, p < 0.001). In Northern Horseshoe, posthoc tests revealed a transient increase in species richness in 2016 (p = 0.04) that eventually returned to the pre-perturbations levels by 2019 (Figure 5A). In Mermaid Cove, the analyses indicated a significant decline in species richness in 2016 (p < 0.001) after the perturbations that did not necessarily recover in the following years (Figure 5B). Furthermore, we used the species accumulation curve to visualise the accumulation of species surveyed as a function of collection effort (transects) (see Supplementary Figure S1).

**Figure 5.**
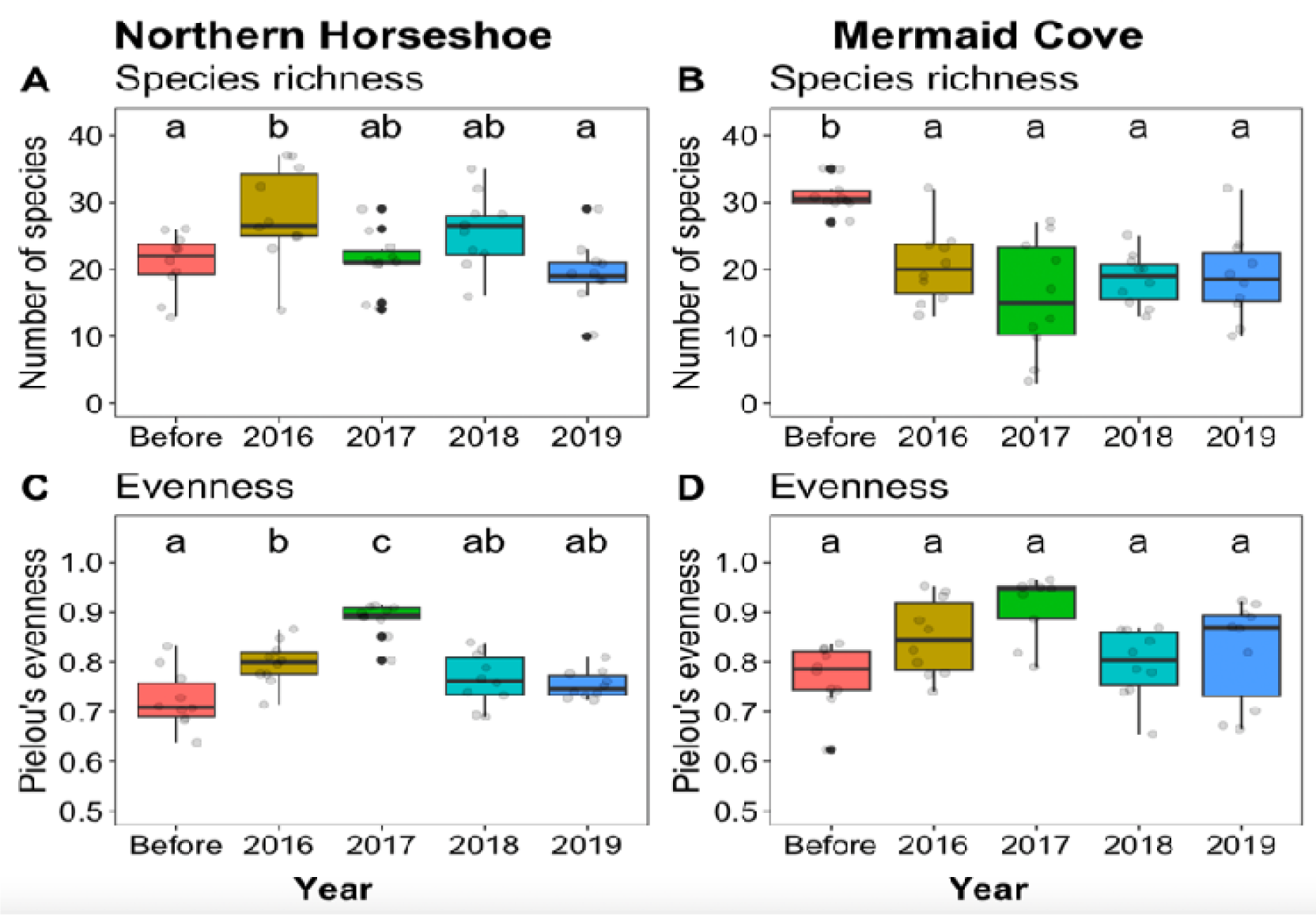
Fish species richness and evenness at Mermaid Cove and Northern Horseshoe. Boxplots of median and interquartile range of (A-B) total number of fish species and (C-D) Pielou’s evenness index. Dissimilar letters above the boxplots indicate statistically significant differences (p < 0.05). The year labelled “Before” corresponds to the data from 2011 for Mermaid Cove and 2014 for Northern Horseshoe. Data from 2022 was not included (see Methods).

#### b. Evenness

Data analyses on Pielou’s evenness index (higher values indicate a more even distribution of species, while lower values towards 0 indicate increased unevenness) showed significant differences across the years (GLS: N =100, X^2^ = 109.871, p < 0.001), but not across locations (GLS: N = 100, X^2^ = 2.0979, p = 0.71). The post hoc analyses revealed significant differences between the years in Northern Horseshoe but not in Mermaid Cove (Figure C&D). For instance, in Northern Horseshoe, there was an increase in Pielou’s index after the perturbations in 2016 (p = 0.04), increasing further in 2017 (p < 0.001) and eventually declining in 2018 and 2019 (p > 0.05) (Figure 5C).

#### c. Composition

Analyses of fish composition showed significant differences across years (PERMANOVA: N = 100, F-value = 8.042, p = 0.001), explaining about 22 % of composition variation. The fish composition was also statistically different across the two locations (PERMANOVA: N = 100, F-value = 8.897, p = 0.001), explaining 6 % of the composition variability. From the visual NMDS plotting of fish composition by year and location (Figure 6), fish composition from before the perturbation clusters together, while the data from the following years can show more overlap among the years and higher dissimilarity (Figure 6).

**Figure 6.**
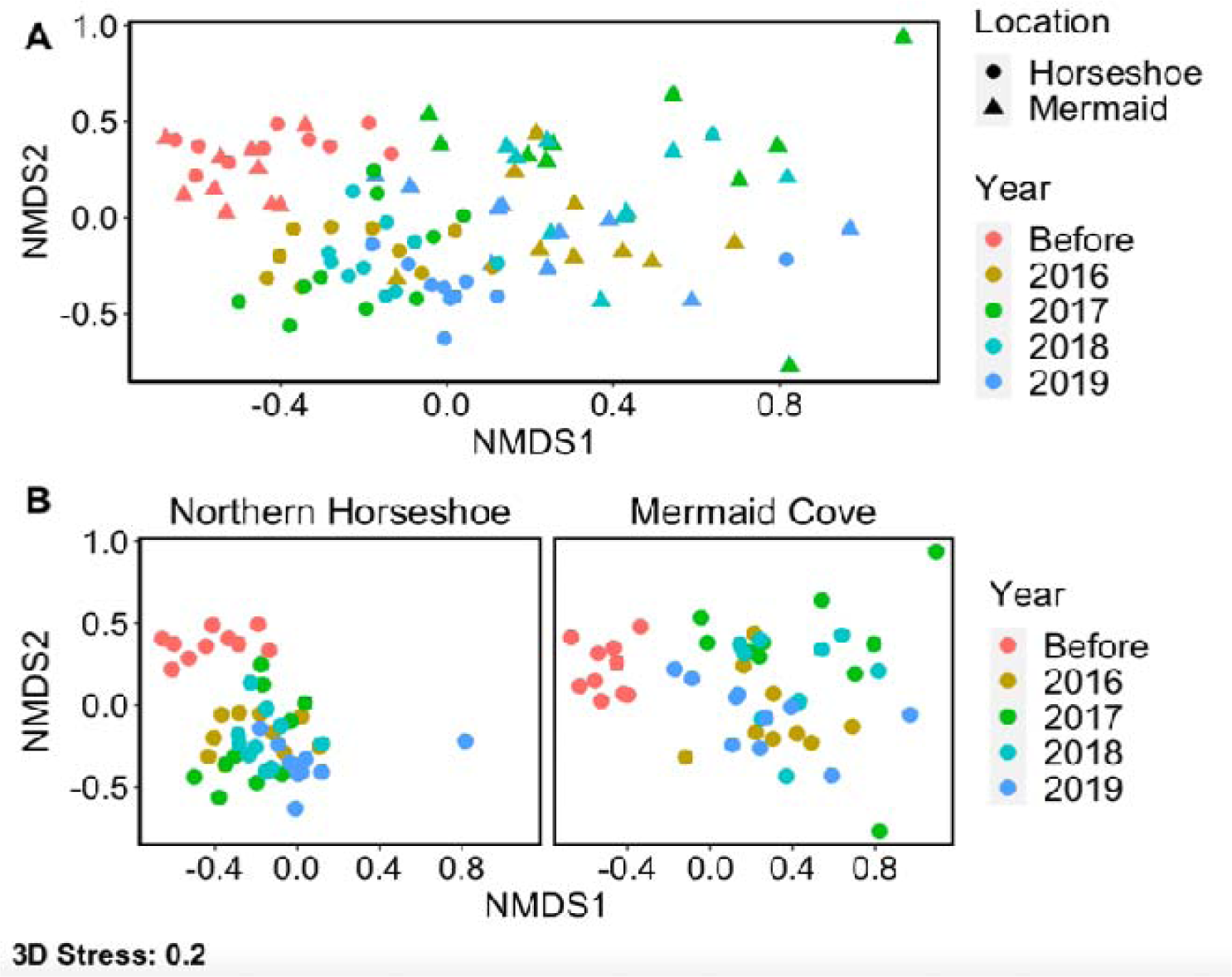
Non-metric multidimensional scaling (NMDS) visualisation of variation in fish composition at Mermaid Cove and Northern Horseshoe. NMDS scatterplot for (A) both locations combined and (B) each location separately. The year labelled “Before” corresponds to the data from 2011 for Mermaid Cove and 2014 for Northern Horseshoe. Each dot represents fish census data from one transect line. Data points that are more similar to one another are ordinated closer together. The further apart the dots, the more dissimilar they are.

## Discussion

The main goal of this study was to gain a comprehensive understanding of how coral reef fish communities recover after repeated environmental disturbances. To achieve this, we used a long-term monitoring approach of the same two locations at Lizard Island between 2011 and 2022. This allowed us to track changes in both fish diversity and abundance following major disturbances.

The results showed significant fluctuation in the recovery of fish communities across the years following the disturbances. Additionally, we observed some differences between the two locations not only after the perturbations but also before, including differences in fish richness and the number of herbivorous fish. The observed pattern differences across the locations after perturbations may be explained by the varying exposure of the two sites to Cyclones Ita and Nathan in 2014 and 2015. For instance, Mermaid Cove, but not Northern Horseshoe, was heavily damaged by the two cyclones (Puotinen et al., 2016; Pizarro et al., 2017). It is, hence, possible that the decrease in fish density and diversity (except for evenness) in Mermaid Cove was caused by this physical disturbance that destroyed the live coral cover and the structural complexity of the location, resulting in fish loss (Wilson et al., 2006; Pratchett et al., 2011). In this case, the coral bleaching had limited impact since the site was already destroyed. While Ceccarelli et al. (2016) did not notice any changes in overall fish density and species richness after cyclone Ita on Lizard Island, even at exposed sites, data collection for that study ended in 2015, and Mermaid Cove was not part of the surveyed locations.

Despite being located in a sheltered area, the Northern Horseshoe reef experienced a significant reduction in fish population by 2017. Although the reef was not physically damaged by the cyclones, it was greatly affected by coral bleaching in 2016. Interestingly, despite the bleaching, there was an increase in species richness and evenness, but this did not translate to a rise in the abundance of fish in fish census data from 2016. A possible explanation is that the data was collected only a few months after the onset of the bleaching (Triki et al. 2018) when the corals were still alive but bleached (survival up to six months without zooxanthella (McCook, 2002; Pratchett et al., 2008)), which might have increased the heterogeneity of the habitat making it easier for the observer to detect and count cryptic fish species. This remains difficult to confirm without information about the benthic habitat structure and benthic communities to clearly assess how they may contribute to fish census data collection. Due to the erosion and death of coral skeletons, the reef’s structural complexity decreased over time, which may have resulted in the significant decline in fish abundance in 2017. It is also possible that the shallow depth of the site (< 5m) accelerated the erosion (Sheppard et al., 2002). The increase in evenness in 2017 and the fact that the fish density of all functional groups decreased, except corallivores, possibly due to their general low density, indicated that the fish loss was even and resulted in similar densities across species. It suggests that disturbances affecting corals can affect all reef fish, regardless of their direct dependence on them (Triki & Bshary, 2019).

Fish census data from 2022 showed a remarkable recovery in fish abundance five years after the last major disturbance event. In both locations, fish density reached similar levels as before the perturbations. This is in line with previous findings at Lizard Island across 14 reef sites showing a significant recovery in fish abundance between the years 2017 and 2020 (Richardson et al. 2021). This is possibly linked to evidence suggesting coral reef recovery at Lizard Island (see Tebbett et al., 2022). Researchers have estimated that corals might take about 6 to 15 years to recover (Morri et al., 2015; Ortiz et al., 2018; Osborne et al., 2017; Tebbett et al., 2022). However, some coral reefs might take longer to recover (Jaroensutasinee et al., 2020), and it can be challenging to recover when the time interval between extreme disturbances is short (Hughes et al. 2018).

In Northern Horseshoe, fish populations recovered quicker than in Mermaid Cove. The total abundance and functional group abundances, as well as richness and evenness, returned to pre-disruption levels by 2018 in Northern Horseshoe and by 2022 in Mermaid Cove. The recovery speed differences might be again explained by the fact that Mermaid Cove suffered severe physical damage that would take longer to recover compared to bleached corals. Also, bleached corals can experience a shift in coral-algal growth where algae start to proliferate while corals undergo erosion, like in the Caribbean (e.g. Hughes, 1994). While the shift from coral to algae may explain the increase in herbivore population that feeds on algae, it is not the only reason behind the rise in omnivore and carnivore densities. It is possible that omnivores adapted their dietary habits to include more plant material as a result of the shift, while the surge in fish prey populations may have contributed to the increase in carnivore density.

We hypothesise that the recovery of coral is the main factor that has led to the recovery of fish abundance between 2019 and 2022. However, we cannot overlook the potential impact of the COVID-19 lockdown. Human activities have been known to have a significant impact on the dynamic of reefs (Schipper et al., 2008). The pandemic lockdown has significantly reduced human activities from 2020 to 2022, which has already shown a positive effect on the environment and wild animals (Arora et al., 2020; Bertucci et al., 2021). Therefore, it is possible that the prolonged lockdown at Lizard Island led to a general increase in fish density due to the large reduction in human activities. Recent studies have shown a significant increase in coral reef fish densities during the lockdown period (Feeney et al., 2022; Bertucci et al., 2023). However, it is also possible that the lockdown affected the fish assemblage, which resulted in higher numbers during surveys, but this could lead to a decline in fish numbers once the lockdown is lifted and human activities resume (Lecchini et al., 2021). In our case, continuous surveys in the years after 2022 will help to clarify this issue.

In conclusion, our research presents a case study that sheds light on how fish density and diversity fluctuate in response to repeated disturbances. This study offers new insights into the recovery of fish populations in terms of density, richness, evenness and composition in areas that have been disturbed. It also underscores the complexity and variability of ecosystem responses to disturbances and highlights the importance of long-term monitoring surveys.

## Ethics statement

The University of Queensland Animal Ethics Committee (AEC) approved the study under permits: CA 2017-05-1063, CA 2019-06-1285 and CA 2022-04-1601.

## Data availability

The data and code will be available upon peer review publication.

## Conflict of Interest

All authors declare that they have no conflict of interest.

## Acknowledgements

We thank the LIRS directors and staff for their support and friendship. We sincerely thank Sergio Rasmann and Radu Slobodeanu for their assistance and advice on the statistical analyses. Financial support was from the Swiss National Science Foundation (grant numbers 310030B_173334/1 to RB and PZ00P3_209020 to ZT).

## Author contribution

ZT and RB designed the study. RB, ZT and LP collected the data. SL ran the statistical analyses, generated the figures, and wrote the manuscript with input from RB and ZT. All authors gave final approval for publication and agreed to be held accountable for the work performed therein. A preprint version of this article is available on bioRxiv (Levy et al. 2024).

## Electronic Supplementary Material

**Figure S1.**
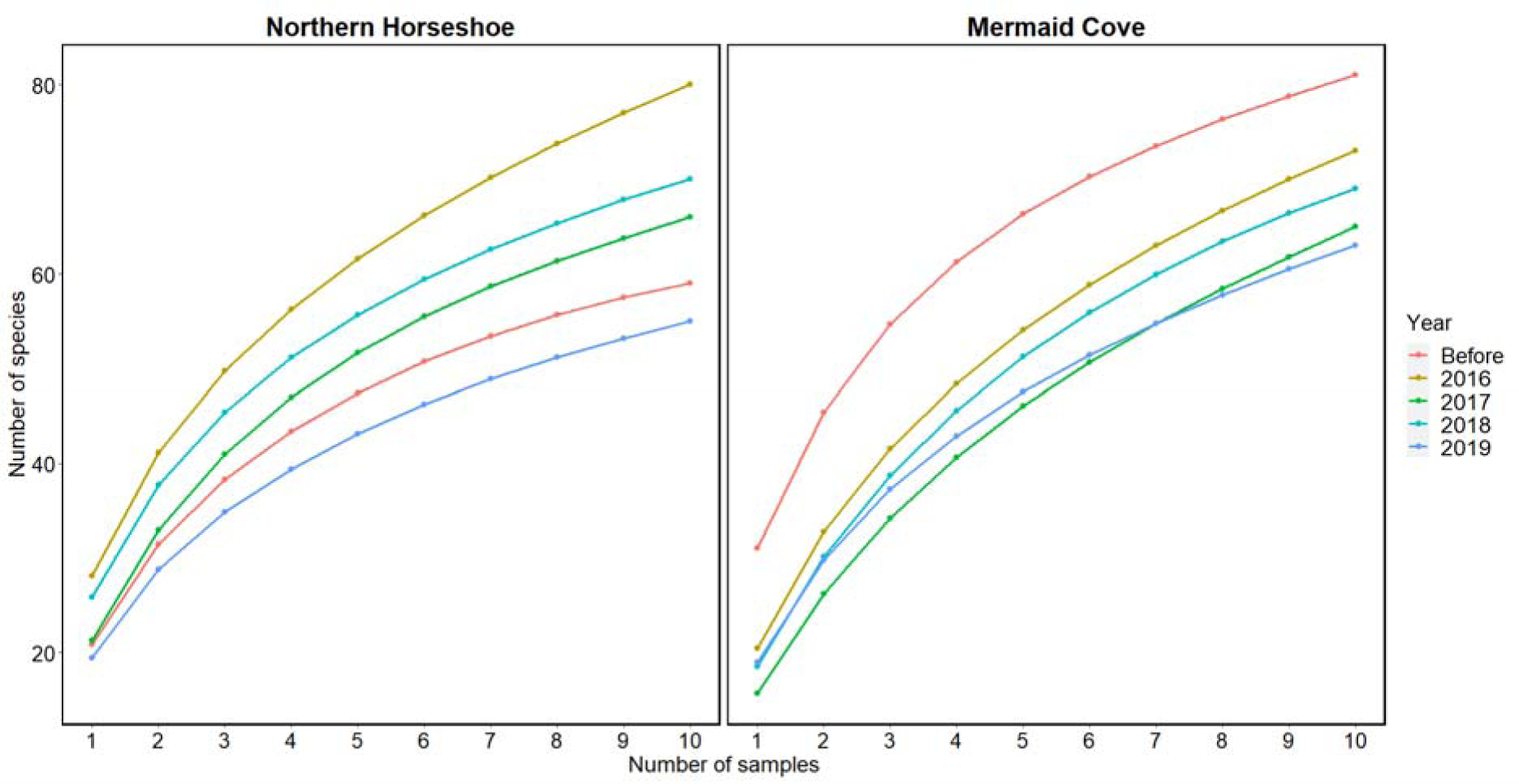
Species accumulation curves at Mermaid Cove and Northern Horseshoe between 2011 and 2019. Each curve represents the average of 10,000 permutations of sampling order. The year labelled “Before” corresponds to the data collected in 2011 from Mermaid Cove and 2014 from Northern Horseshoe before the disturbances. The x-axis corresponds to the number of sampled transects (10 transects per location and period of data collection).

**Table S1.**
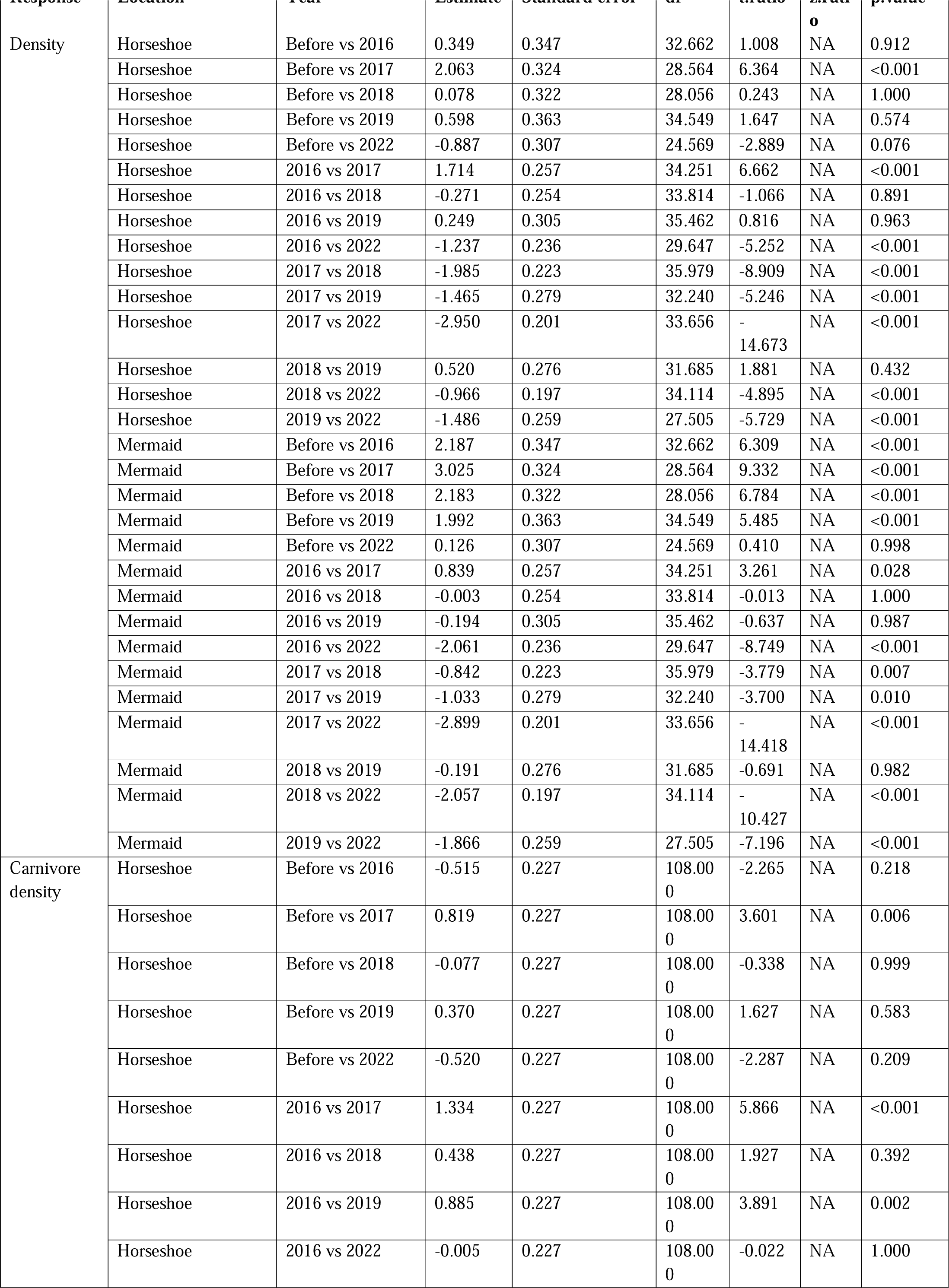
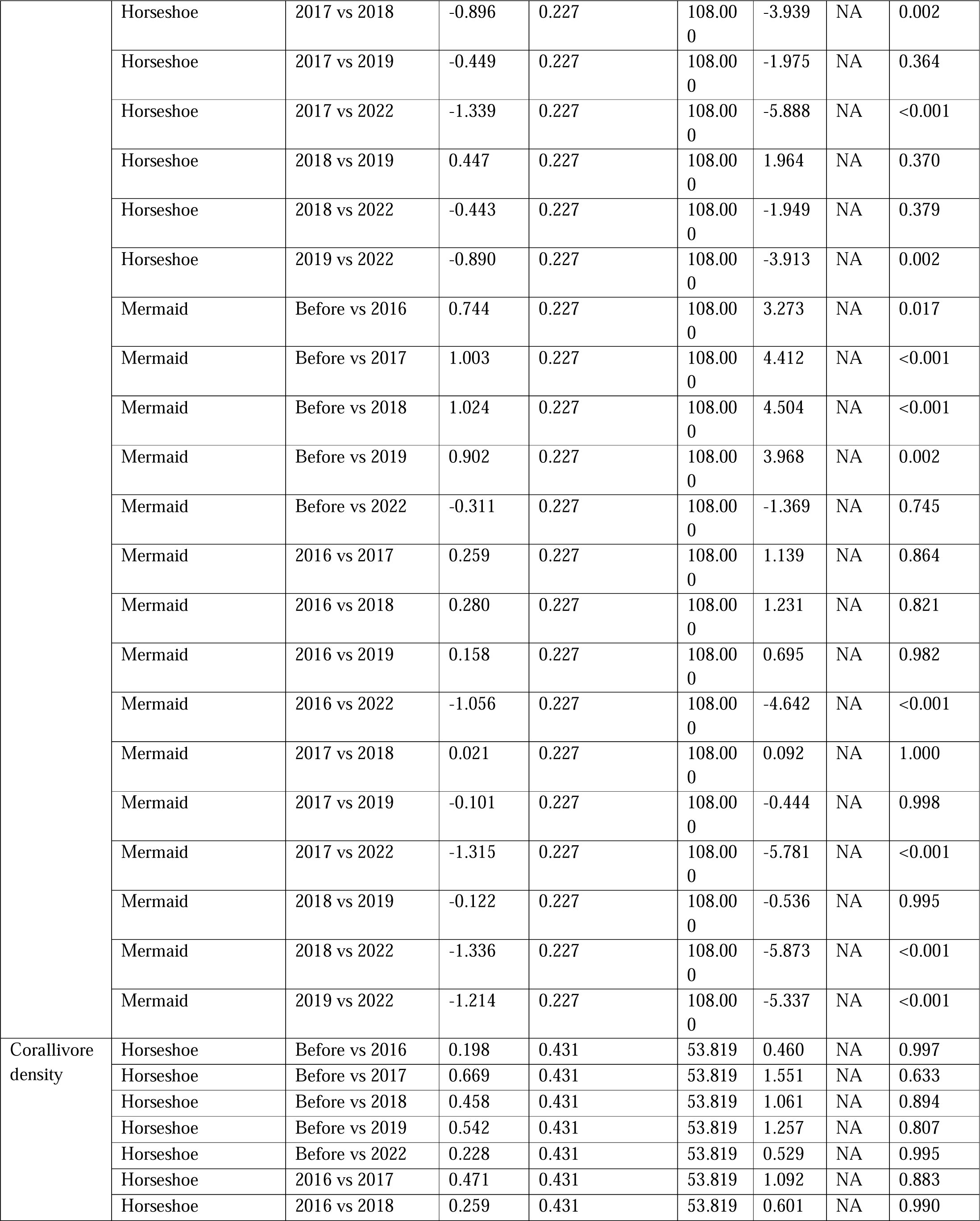
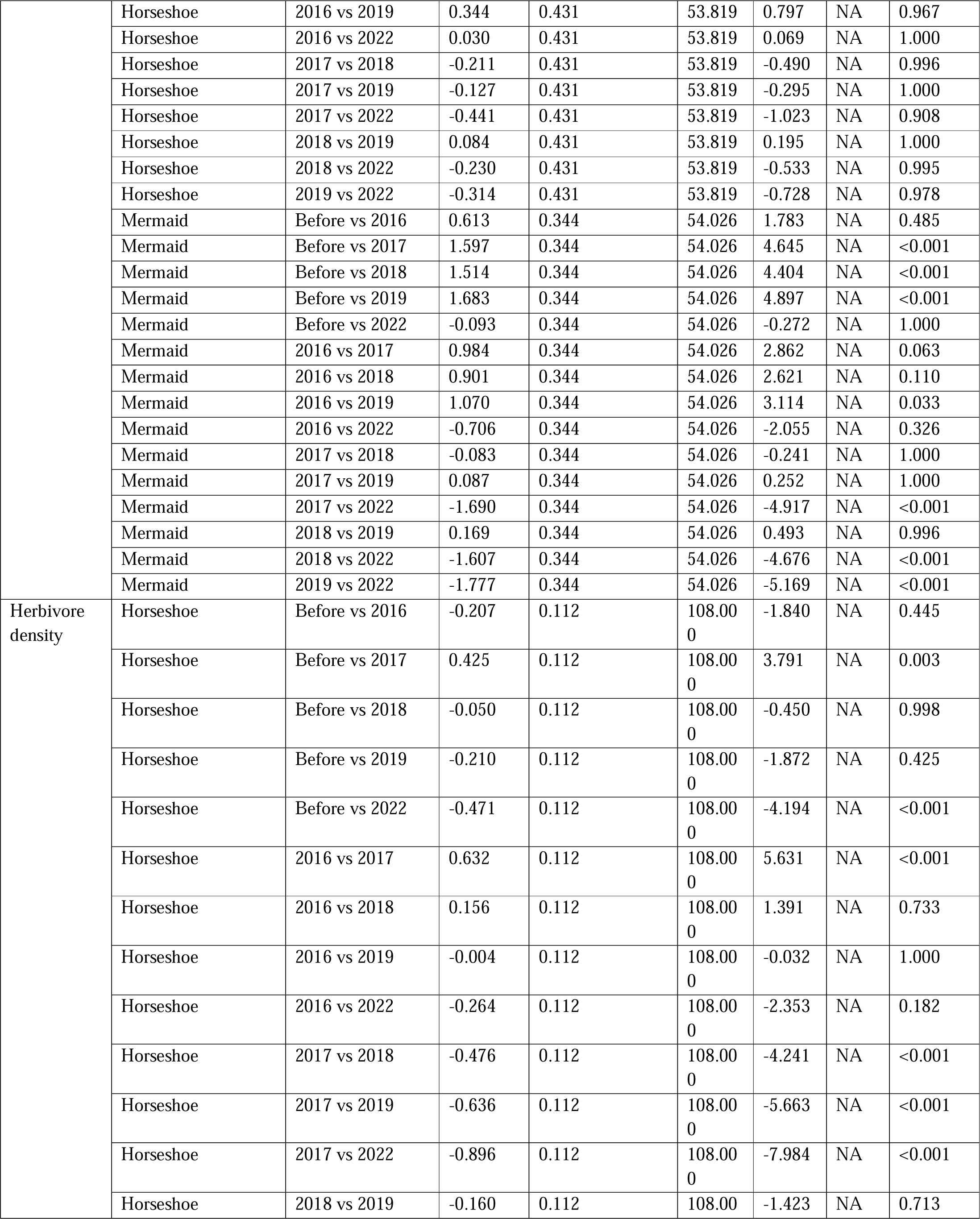
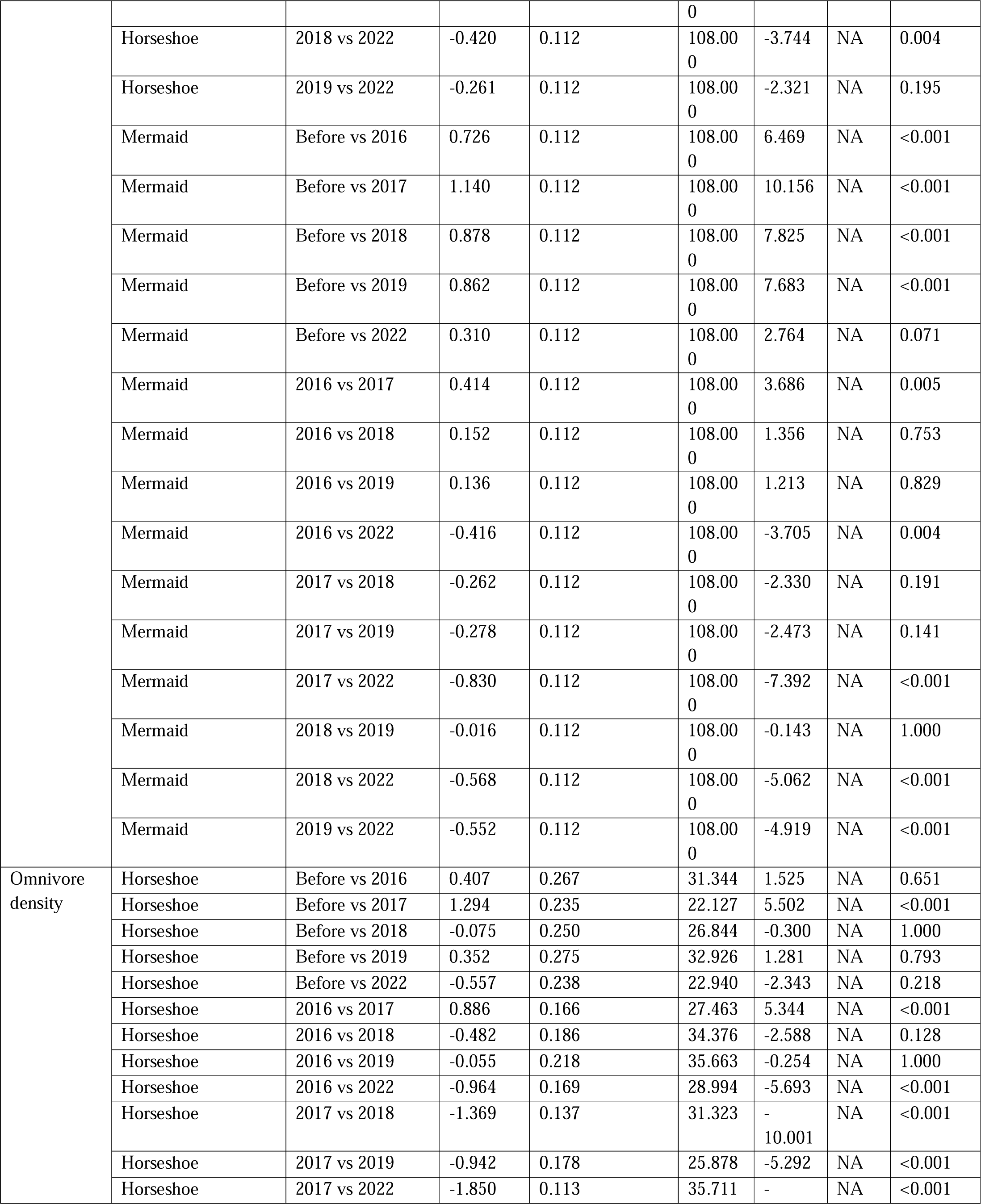
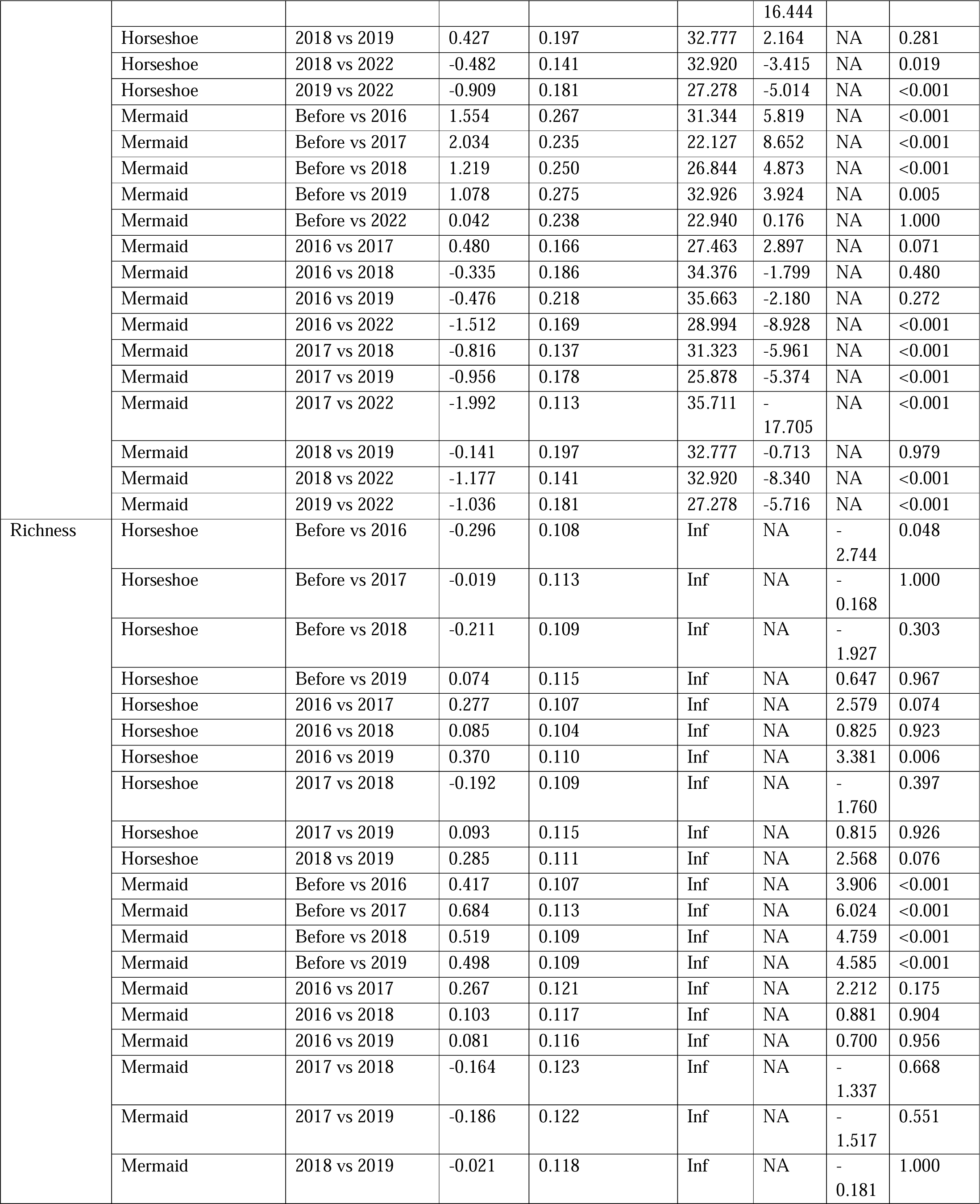
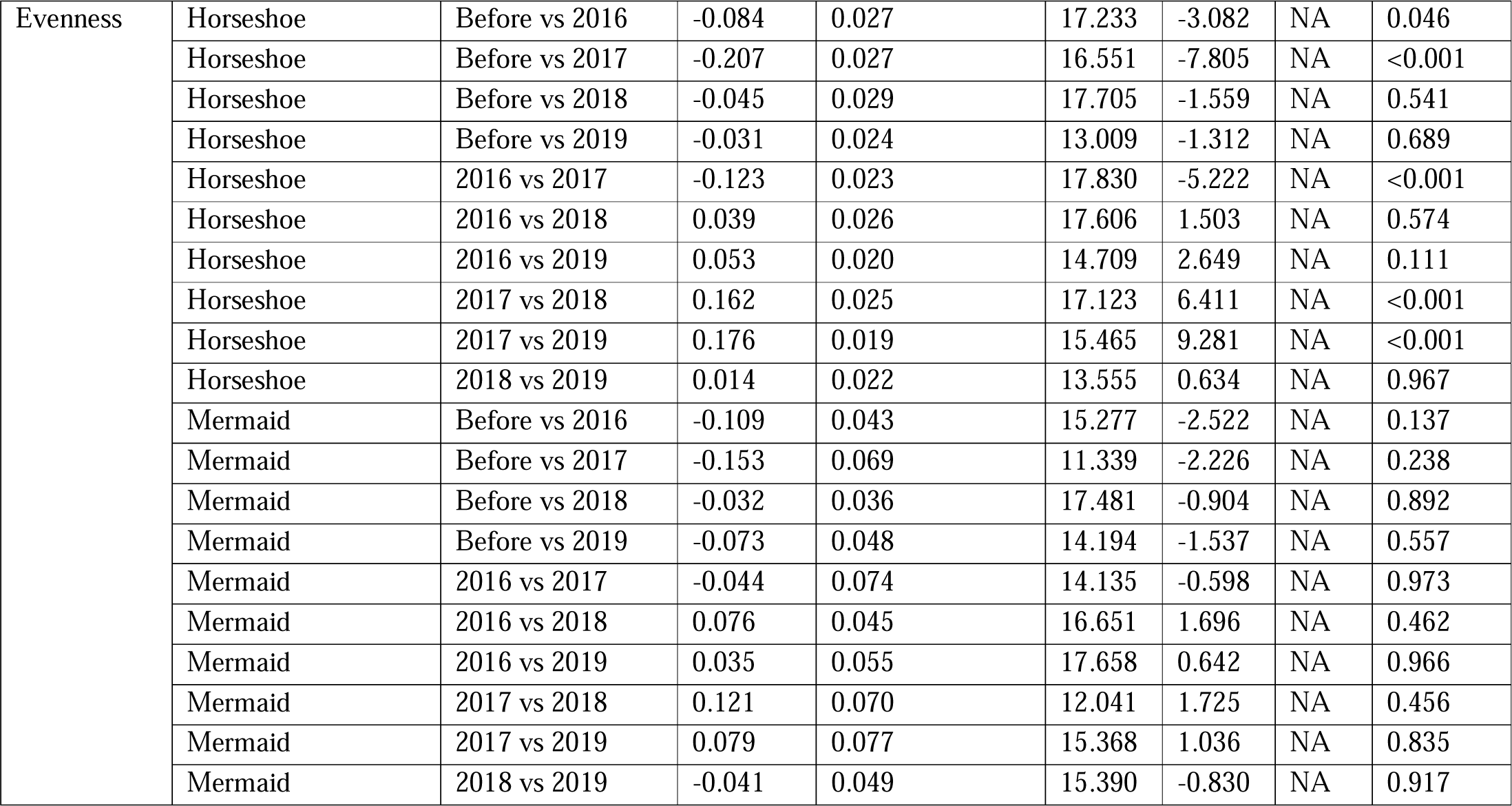
Post-hoc analyses’ outcomes for the tested response variables. The location labelled “Horseshoe” corresponds to Northern Horseshoe and “Mermaid” to Mermaid Cove. The year labelled “Before” corresponds to the 2011 data from Mermaid Cove and 2014 from Northern Horseshoe.

## Notes

### Competing Interest Statement

The authors have declared no competing interest.

